# Axonal pathology differentially affects human Purkinje cell subtypes in the essential tremor cerebellum

**DOI:** 10.1101/2025.01.26.633063

**Authors:** James Widner, Phyllis L. Faust, Elan D. Louis, Hirofumi Fujita

## Abstract

The cerebellar cortex is organized into discrete regions populated by molecularly distinct Purkinje cells (PCs), the sole cortical output neurons. While studies in animal models have shown that PC subtypes differ in their vulnerability to disease, our understanding of human PC subtype and vulnerability remains limited. Here, we demonstrate that human cerebellar regions specialized for motor vs cognitive functions (lobule HV vs Crus I) contain distinct PC populations characterized by specific molecular and anatomical features, which show selective vulnerability in essential tremor (ET), a cerebellar degenerative disorder. Using a known PC subtype marker, neurofilament heavy chain (NEFH), we found that motor lobule HV contains PCs with high NEFH expression, while cognitive lobule Crus I contains PCs with low NEFH expression in post-mortem samples from healthy controls. In the same cerebella, PC axons in lobule HV were 2.2-fold thicker than those in Crus I. Across lobules, axon caliber positively correlated with NEFH expression. In ET cerebella, we identified motor lobule-specific PC axon pathology with a 1.5-fold reduction in caliber and increased axon variability in lobule HV, while Crus I axons were unaffected. Tremor severity and duration in ET correlated with axon diameter variability selectively in lobule HV PCs. Given that axonal caliber is a major determinant of neural signaling capacity, our results (1) suggest that disrupted cerebellar corticonuclear signaling is occurring in ET, (2) provide evidence of region-specific PC subtypes in the human cerebellum and offer insight into how selective PC vulnerability may contribute to the pathophysiology of cerebellar degeneration.

**Significance Statement:** The cerebellar cortex has a uniform laminar architecture but contains heterogeneous cell types. Purkinje cells (PCs), the sole output cells of the cerebellar cortex, include subtypes whose lobular distribution is thought to underlie functional segmentation and patterned degeneration of the cerebellum in animal models. However, human PC subtypes and their disease vulnerability remain unknown. Here, we establish the existence of human PC subtypes that appear conserved across mammalian species using both marker expression and axonal thickness. Consistent with its phenotype, we demonstrate differential PC vulnerability to degeneration in essential tremor cerebella, where motor function- mediating PC subtypes display significant axonal thinning, while axons of cognitive function-mediating PC subtypes are spared. These findings advance Purkinje cell type-oriented research in cerebellar disorders.

## Introduction

The cerebellar cortex has been traditionally viewed as a homogeneous structure with a uniform cellular architecture. However, studies have begun to identify extensive heterogeneity within it, revealing topographical differences in terms of neuronal subtypes, connectivity patterns, functional roles, and manifestations in neurodegenerative diseases (1–6). Of particular interest is the broad functional dichotomization of the cerebellar cortex into motor and non-motor regions, as shown in rodents, non-human primates, and humans (7–10). Despite these advances, a critical gap remains in our understanding of the human cerebellum, specifically in the characterization of Purkinje cell (PC) subtypes and their differential vulnerability to disease.

PCs are the sole output neurons of the cerebellar cortex, transmitting cerebellar cortical signals to the cerebellar nuclei. Animal studies have identified that molecularly and topographically distinct PC subsets are integrated into functionally distinct output circuits (3–5, 11, 12). Importantly, these PC subtypes can also be distinguished by their axonal caliber, a critical structural characteristic that supports precise, well-timed signal transmission (13–18). Here, we hypothesize that in the human cerebellum, differences in PC axon caliber could distinguish PC subsets that are associated with different output circuits (i.e. motor or non-motor regions (19)) and that these subsets might exhibit varying responses to disease.

To understand human PC subtypes from a molecular, anatomical, and pathophysiological perspective, we compared distinct PC populations from post-mortem control and essential tremor (ET) cerebella. ET is among the most common neurological diseases and the most common cerebellar degenerative disease (20); its prevalence is extraordinarily high, with an estimated 7 million affected people in the United States, representing 2.2% of the entire US population (21, 22). Patients with ET present with motor-dominant symptoms, typically an action tremor of the upper limbs, but a host of other cerebellar features may occur as well (intention tremor, ataxia, problems with motor timing) (23, 24). This is paralleled by cerebellar neurodegeneration and prominent pathological changes in the PCs (25). Thus, pathological investigation of the ET cerebellum could allow targeted examination of PCs across dichotomized cerebellar functional domains.

In this study, we first identified molecularly distinct PC subtype populations, which parallel those of rodent species using an established marker of PC subtype, neurofilament heavy chain (NEFH). We then 3D-reconstructed 1586 PC axon segments in control and ET cases from these lobules (**Fig. 1**) to examine differences between HV and Crus I PCs in healthy controls and ET patients. The results show axonal structural fidelity as a potential pathophysiological mechanism underlying cerebellar motor symptoms in ET.

**Figure 1.**
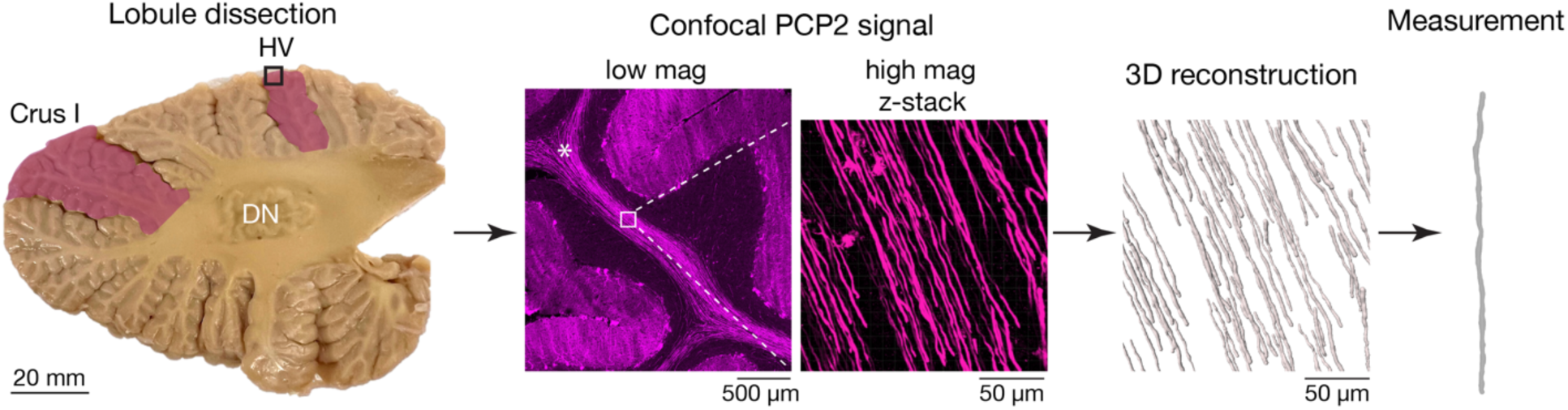
**3D-reconstruction and measurements of human PC axons.** Analytic flow of single axon measurement in this study. Left, Parasagittal slice of lateral human cerebellum (∼1.5-2.5 cm from midline, at the dentate nucleus (DN), in which hemispheric lobule V (HV) and lobule Crus I are stereoscopically identified (pink shading). Middle, apical regions of the tissue slice (black rectangle in left) are dissected and sectioned. Purkinje cell (PC) bodies and axons are visualized via fluorescent immunostaining for PCP2. High-magnification z-stack image is acquired from a white rectangular area slightly basal to the apical most convergence point of the cerebellar folia (indicated by white asterisk). Right, z-stack images are processed in Imaris software for three-dimensional reconstruction of individual PC axons. Subsequently, axonal measurements are performed on these reconstructed 3D models.

## Results

### Demographic and clinical features

Demographic and clinical features are shown in ET cases and controls (**Table 1**). On average, ET cases had tremor for 54 years and average total tremor score was 22, indicating the presence of moderate to severe kinetic tremor on multiple tasks (e.g., pouring, drinking, using a spoon, finger-nose-finger maneuver, and spiral drawing). There were no significant case-control differences in post-mortem interval, total brain weight, Braak Alzheimer’s Disease stage, or CERAD score, but age between groups differed slightly (**Table 1**). We later controlled for this age difference by selecting 7 of the 10 ET cases that were most similar to each control in age to construct an age-matched sample of 7 ET cases and 5 controls. This selection was made blinded to any clinical information other than age and any other results from our analyses (**Table S1**). Mean time (± SD) from brain sample acquisition to analysis was similar for ET and controls (ET = 4.0 ± 2.3 years, range 1.2 – 8.8 years; control = 4.2 ± 2.6 years, range = 0.4 – 8.6 years; *p* = 0.57, Mann Whitney test); 80% of both ET and control samples were acquired within the past ∼3.2 years. On average, ET individuals were 5 years older than controls, a difference that did not reach statistical significance (**Table 1**). Nevertheless, we later controlled for age by selecting 7 ET cases and 5 controls that were most similar in age, thereby constructing an age-matched sample. This selection was made blinded to any clinical information other than age and any other results from our analyses (**Table S1**).

**Table 1.** Clinical and pathological characteristics of samples.

|  | Control | ET | Comparison |
| --- | --- | --- | --- |
| N | 6 | 10 |  |
| Age (years) | 83 ± 8 | 88 ± 5 | p = 0.09<sup>a</sup> |
| Female sex | 2 (33%) | 7 (70%) | p = 0.30<sup>b</sup> |
| Tremor age of onset (years) | NA | 34 ± 22 |  |
| Tremor duration (years) | NA | 54 ± 24 |  |
| Head tremor on examination | NA | 5 (50%) |  |
| Voice tremor on examination | NA | 5 (50%) |  |
| Only arm tremor on examination | NA | 4 (40%) |  |
| Total tremor score | NA | 22 ± 6 |  |
| PMI (hours)* | 28 ± 24 | 32 ± 16 | p = 0.42<sup>a</sup> |
| Total brain weight (g) | 1,269 ± 88 | 1,189 ± 130 | p = 0.18<sup>a</sup> |
| Braak AD stage |  |  | p > 0.99<sup>b</sup> |
| 0-4 | 5 (83%) | 7 (70%) |  |
| 5 or 6 | 1 (17%) | 3 (30%) |  |
| CERAD score |  |  | p = 0.89<sup>b</sup> |
| 0 | 2 (33%) | 3 (30%) |  |
| A | 1 (17%) | 2 (20%) |  |
| B | 1 (17%) | 4 (40%) |  |
| C | 2 (33%) | 1 (10%) |  |

All values are mean ± standard deviation or number (percentage).
\*PMIs (time between patient death and brain samples freezing) are unavailable for 1 control sample.
<sup>a</sup>Student's t-test
<sup>b</sup>Fisher's exact test
Abbreviations: AD, Alzheimer's disease; CERAD, Consortium to Establish a Registry for Alzheimer's Disease; PMI, post-mortem interval.

### Human PC subtypes are distinguished by neurofilament expression and axon caliber

To distinguish PCs based on molecular expression, we performed immunostaining in six controls for neurofilament heavy chain (NEFH), a marker of PC subtype (26). In the stained cerebellar sections, we measured the mean fluorescence intensity within each PC somata in lobules HV and Crus I (n = 93 and 59, respectively). PCs in HV were strongly immunopositive relative to those from Crus I PCs (**Figs. 2A,B**). In a paired analysis, every control subject exhibited greater NEFH PC expression in lobule HV compared to Crus I (**Fig. 2C**; 74 ± 10 vs 42 ± 18 a.u.; paired t-test; *p* = 0.0072).

**Figure 2.**
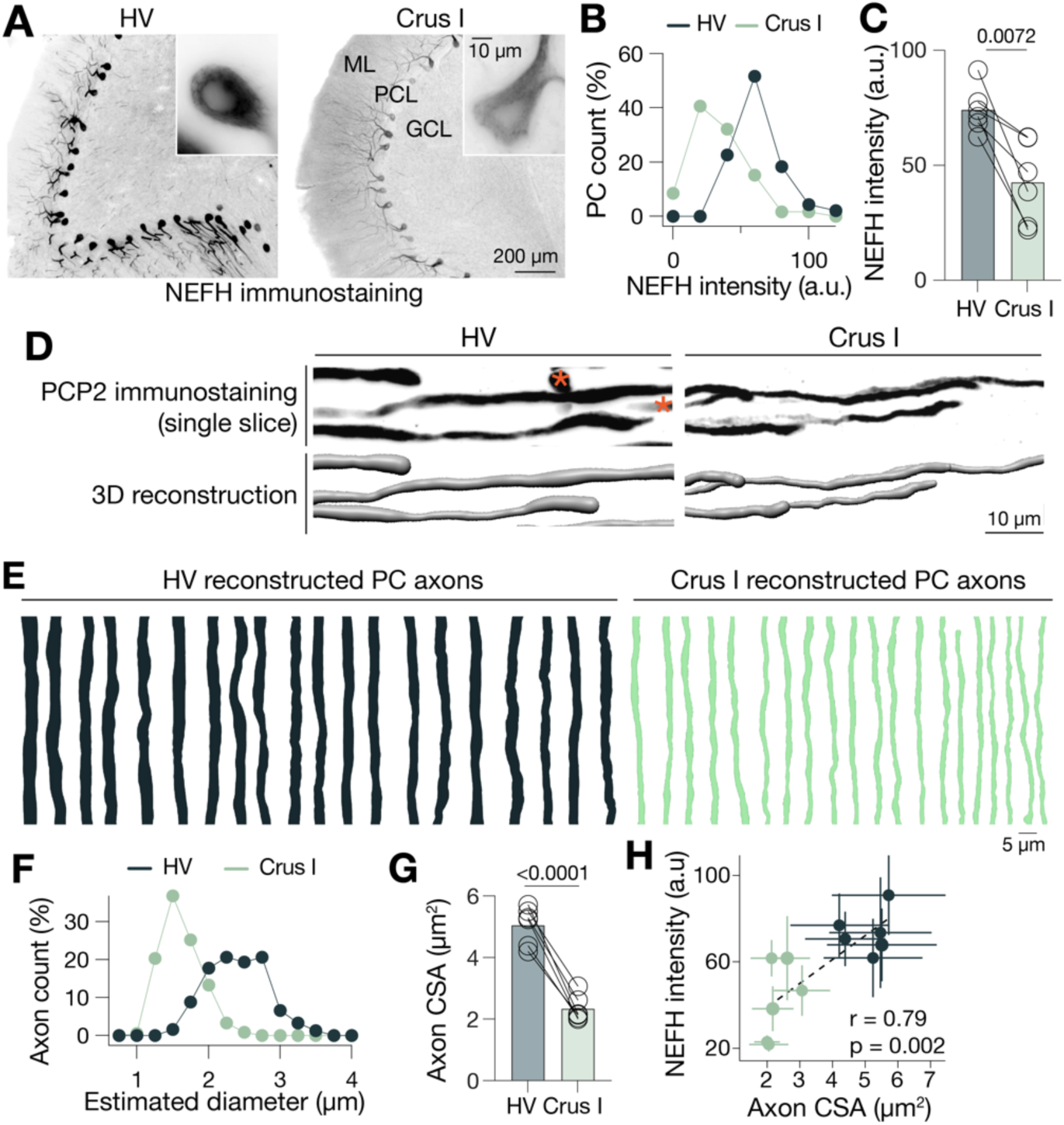
PC subtypes across lobules HV and Crus I in healthy controls. **A.** Fluorescent immunostaining of the human cerebellar cortex for neurofilament heavy chain (NEFH). Images of lobules HV and Crus I were taken from the same subject using identical experimental conditions and exposure time. Darker signal indicates greater immunoreactivity. Note more intense somatic signals in HV PCs (left) compared to those in Crus I (right). Insets show representative PC somata in these regions. **B.** Histogram of PC counts in lobules HV and Crus I sorted by NEFH intensity (n = 6 subjects; 93 vs 59 cells, respectively; bin size is 20 a.u.). Vertical axis represents the count in each bin as a fraction of the total PC count. **C.** Paired analysis of NEFH intensity within samples. Average values of measured intensity in each subject are plotted. Every subject exhibited greater PC NEFH intensity in HV PCs versus that in Crus I (n = 6; paired t-test, t(5) = 4.367, *p* = 0.0072). Compared across individuals, NEFH immunoreactivity was ∼1.7-fold greater in HV (74 ± 10 vs 42 ± 18 a.u., mean ± SD of n = 6 data points). **D.** Example of PC axon 3D reconstruction from confocal image. High magnification confocal image (top) is imported into Imaris for 3D reconstruction (bottom) and quantification. Examples from both HV and Crus I are shown. Orange asterisks indicate axons that were not reconstructed. **E.** Examples of 3D-reconstructed PC axons from lobules HV and Crus I. Forty example axons are shown. **F.** Histogram of estimated axonal diameter of PCs in lobules HV and Crus I. 305 vs 370 PC axons from HV and Crus I, respectively, from 6 healthy controls; bin size is 0.25 µm). Axon diameter was estimated from cross-sectional area (CSA), assuming circularity. **G.** Paired analysis of PC axon CSA from lobules HV and Crus I within samples. Average values of CSA in each subject are shown with Tukey box plot. Lobule HV PC axons were significantly thicker than those in lobule Crus I (n = 6; paired t-test, t(5) = 11.76, *p* < 0.0001). Compared across individuals, PC axon CSA was ∼2.2-fold greater in HV (5.1 ± 0.6 vs 2.3 ± 0.4 µm^2^, mean ± SD of n = 6 data points). **H.** Distinct population of PCs in lobules HV and Crus I identified by NEFH intensity and axonal CSA. Each point is an averaged value ± SD of the measures in either HV (dark green) or Crus I (light green). Twelve points from six subjects are shown. Linear correlation in this plot (Pearson’s r = 0.79, p = 0.002) is consistent with the known association between neurofilament expression and axon caliber found in animal models (66). Abbreviations: GCL, granule cell layer; HV, hemispheric lobule V; ML, molecular layer; PCL, Purkinje cell layer.

To examine whether human PCs can be distinguished by axonal caliber, we selectively visualized PC axons in the white matter by immunostaining for a specific pan-PC marker, PCP2 (**Fig. 1**). The labeling revealed robustly stained PC axons at low magnification (**Fig.1**), which exhibited regional micro-variations in thickness when viewed at high magnification (**Fig. 2D**). To represent axonal caliber, we thus calculated the cross-sectional area (CSA) by dividing axon volume by axon length. To do so, we 3D-reconstructed the labeled PC axons randomly from the viewfields of confocal z-stacks, and the volume and length were measured in each of the reconstructed axonal segments (**Figs. 1, 2D,E**; Methods). 305 and 370 axons were quantified in HV and Crus I, respectively. Comparison between the obtained CSAs demonstrated a remarkable difference between the CSA of the axons in lobules HV and Crus I (**Figs. 2F,G**; 5.1 ± 0.6 vs 2.3 ± 0.4 µm^2^, respectively; paired t-test; *p* < 0.0001). HV axons showed significantly greater variability in their thickness than that of Cr1 (measured as CV of CSAs, 31 ± 3 vs 27 ± 4 %; paired t-test, t(5) = 2.623; *p* = 0.047). Further, we identified a strong positive correlation between PC axon caliber and NEFH intensity in lobule HV and Crus I PCs in controls (**Fig. 2H**; r = 0.79, *p* = 0.002).

Taken together, we identified distinct populations of human PC subtypes based on NEFH expression and axonal caliber. The observed pattern, in which NEFH-high PCs in HV had thicker axons than NEFH-low PCs in Crus I, parallels what is observed in PC subtypes of animal models (16, 26).

### PC subtypes exhibit selective vulnerability in ET

To test if the PCs in lobules HV vs Crus I demonstrate disease-specific vulnerability, we examined PC axons in the patients with essential tremor (ET), a degenerative disorder characterized primarily by action tremor. Prior to analyzing PC axons in ET cerebella, we wanted to ensure that PCP2 could still be used as a reliable marker to stain all PCs in the ET cerebellum, as PCs could change gene expression under disease conditions (27, 28). To test this, we quantified the fraction of PCP2-labeled somata among the Nissl-identified PC somata. We confirmed that 99.1% of PC somata were labeled with PCP2 in both ET and control cerebella (measured in HV, n = 4 samples each; n = 993 vs 424 cells for control and ET, respectively) (**Figs. 3A,B**). This result indicated that PCP2 labeled close to the entire population of PCs both in ET and control and thus could be used as a proxy for CSA measurements in ET PCs.

**Figure 3.**
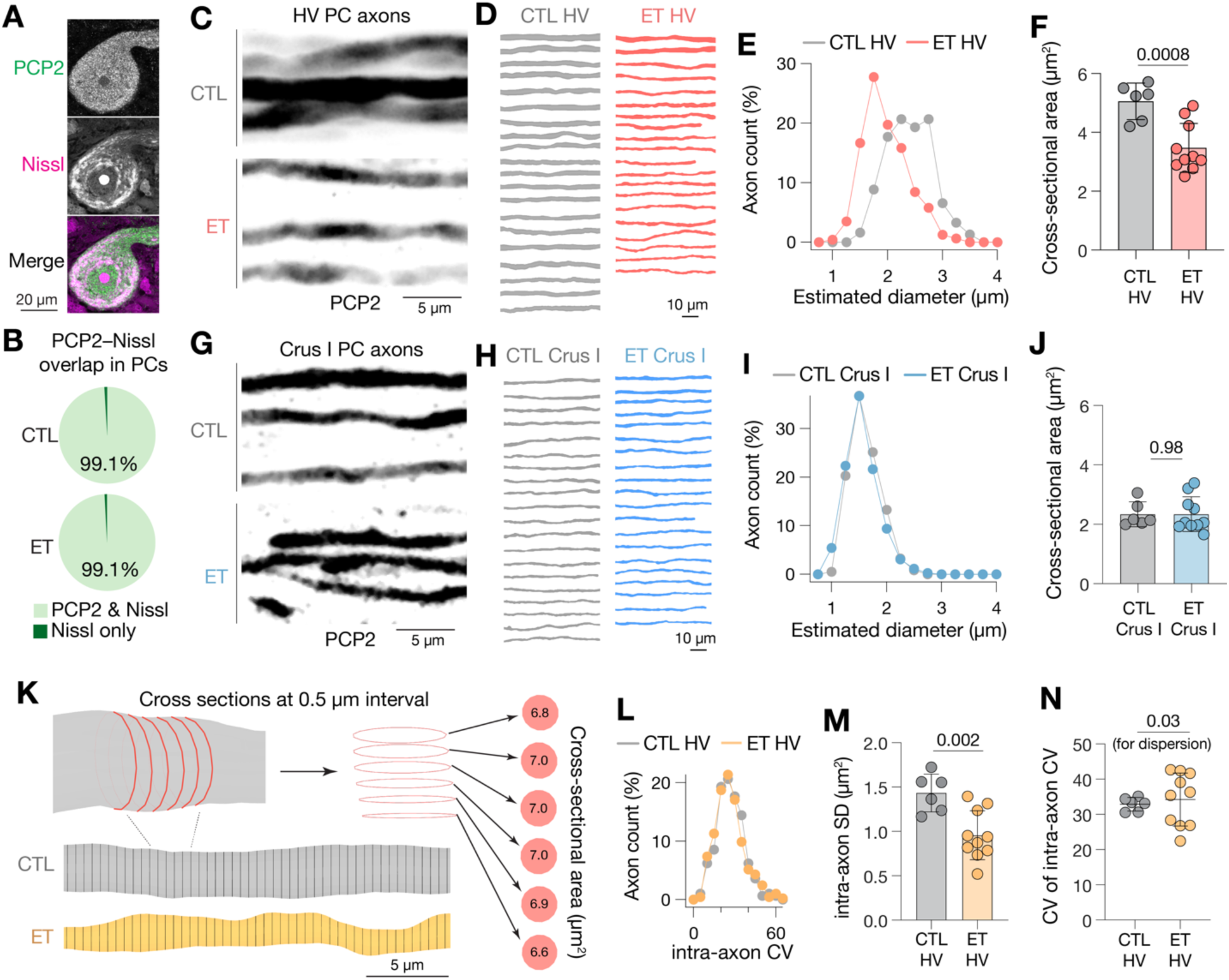
Purkinje cell axons in the ET cerebellum display degenerative morphology in hemispheric lobule V, but not in Crus I. **A.** Immunostaining of ET PC for PCP2. Example result of a PC somata identified with Nissl counterstaining is shown. **B.** Overlap between Nissl and PCP2 immunostaining in PC somata quantified in lobule HV PC layer from control and ET cerebella (n = 4 samples each; n = 993 vs 424 cells for control and ET, respectively; 99.1% vs 99.1% PCP2-Nissl overlap). **C.** Representative high magnification confocal image of lobule HV PC axons in control (top) and ET (bottom) cerebellar white matter. **D.** Examples of 3D-reconstructed HV PC axons from control and ET cerebellar white matter. Twenty examples from each group are shown. **E.** Histogram of reconstructed HV PC axon diameter in control and ET. The diameter derives from cross-sectional area assuming circularity. Left shift of the distribution of ET axons illustrates the general thinning of lobule HV PC axons (n = 6 vs 10 subjects; n = 305 vs 486 axons; bin size is 0.25 µm; control data are also presented in Fig. 2F). **F.** Case comparison of HV PC axon CSA between control and ET. Each point corresponds to an averaged value of all the measured PC cross-sectional areas (µm^2^) in a subject. The HV PC axons were markedly thinner in ET than control (n = 6 vs 10 subjects; unpaired t-test with Welch’s correction, t(13.1)=4.359, *p* = 0.0008; control data are also presented in Fig. 2G). Bar graph indicates mean ± SD (5.1 ± 0.6 vs 3.5 ± 0.8 µm^2^). **G-J**. Analyses on Crus I PC axons, similar to C-F. Histogram of axon diameter does not show shifted distribution between control and ET (I; n = 6 vs 10 subjects; n = 370 vs 425 axons; bin size is 0.25 µm; control data are also presented in Fig. 2F). Case comparison of axon caliber does not reveal statistical distinction between control and ET (J; n = 6 vs 10 subjects; unpaired t-test with Welch’s correction, t(13.4)=0.026, *p* = 0.98; control data are also presented in Fig. 2G). Bar graph indicates mean ± SD (2.3 ± 0.4 vs 2.3 ± 0.6 µm^2^). **K.** Schematics outlining workflow for quantifying within-axon variability of PC axons. 3D- reconstructed axons are segmented orthogonally along their long axis into 0.5 µm-thick sections (bottom, black lines; magnified in top, redline). The rims of these segments are extracted (top, middle) for calculation of their cross-sectional area. Right red disks are example results for surface area calculation (in µm^2^) from the red rims obtained from a control PC axon (gray 3D models, top left and top middle). **L.** Histograms of HV PC axons with various levels of intra-axonal variability. Intra-axonal variability is represented by CV of the all the cross-sectional areas in each axon. These CVs were similar between the axons of the control and the case as a whole (n = 305 vs 486 axons; bin size is 5%), regardless of the difference in the absolute size of the axons. **M.** Case comparison of standard deviation of intra-axon CSA between lobule HV of control and ET. ET HV PC axons show less variation in absolute intra-axon CSA (n = 6 vs 10 subjects; unpaired t-test with Welch’s correction, t(12.9)=3.855, *p* = 0.002). Bar graph indicates mean ± SD (1.4 ± 0.2 vs 1.0 ± 0.3 µm^2^). **N.** Comparison of variability of intra-axon CV in individual subjects between control and ET. Each point is CV of intra-axon CV calculated for each subject. Compared to control group, in which variability of individual axons are as a whole maintained at a similar level, ET group show significant dispersion between subjects (n= 6 vs 10; Ansari-Bradley test; *p* = 0.03). Plots are shown with mean ± SD (33 ± 2 vs 34 ± 7 %).

To examine axon caliber in the ET cerebellum, we 3D reconstructed and quantified the caliber of PC axons as performed in the previous analysis on control cerebellum (**Fig. 2**). Totals of 486 and 425 axons were reconstructed from HV and Crus I PCs, respectively, in 10 ET cerebella. The axon measurements revealed a striking reduction in the caliber of HV PC axons (HV CSA) in ET patients (**Figs. 3C-E**), in which the axonal CSA were, on average, more than 30% smaller in ET than control (**Fig. 3F**; 5.1 ± 0.6 vs 3.5 ± 0.8 µm^2^; unpaired t-test with Welch’s correction; *p* = 0.0008). In contrast, the distribution and average axonal caliber of lobule Crus I PC axons (Cr1 CSA) were indistinguishable in ET and control cerebella (**Figs. 3G-J**; 2.3 ± 0.4 vs 2.3 ± 0.6 µm^2^, respectively; unpaired t-test with Welch’s correction; *p* = 0.98). In the patient group, the variability across HV PC axons (HV CV) was not greater than that of Cr1 (Cr1 CV) (29 ± 6 vs 27 ± 6; paired t-test, t(9) = 1.096, *p* = 0.30), which is influenced by significantly greater dispersion of HV CVs across patients compared to controls (29 ± 6% in ET vs 31 ± 3% in control; Ansari-Bradley test, *p* = 0.03). The selective reduction of caliber in HV PC axons persisted following age matching in a smaller sample (**Table S1**; 5.2 ± 0.5 vs 3.7 ± 0.9 µm^2^; unpaired t-test with Welch’s correction; *p* = 0.0048).

Moreover, all measures of lobule HV PC axon in the present study did not significantly correlate with age, PMI, shelf time, or other changes linked to neurodegeneration (**Table S2**).

To better understand the mode of the thinning observed (e.g. points of focal thinning throughout the axon vs diffuse thinning along the axon), we measured intra-axonal variability, or the extent to which the caliber of an individual axon varies along its length. For this, we split each HV PC axon into small segments with 0.5 µm intervals along their long axes. Intra-axon CV (intraCV) was calculated as a variability of CSAs of these segments (**Fig. 3K**). Overall, intra-axon CVs between ET and control PC axons (n = 305 vs 486) were similarly distributed (**Fig. 3L**). Across-subject comparison also did not reveal differences in averaged intra-axon CVs (intraCV) between ET and control (n = 6 vs 10 subjects; 29 ± 2 vs 28 ± 3 %; unpaired t-test with Welch’s correction; p = 0.66). Notably, however, the absolute size of variability (quantified as the SD of intra-axonal CSAs per patient; intraSD) was significantly decreased in ET axons (**Fig. 3M**; 1.4 ± 0.2 µm² in controls vs 1.0 ± 0.3 µm² in ET; n = 6 vs 10 subjects; unpaired t-test with Welch’s correction, *p* = 0.0022). This represents a 1.4-fold decrease in the dynamic range of intra-axonal size. In addition, we found that the variability of intra-axon CV values (CV of intra-axon CV; intraCVCV) was tightly regulated in controls (33 ± 2%; n = 6) but showed significantly greater dispersion in ET patients (34 ± 7%; n = 10; Ansari-Bradley test, *p* = 0.03; **Fig. 3N**). Together, these results suggest that HV PC axons in ET patients exhibit decreased and dysregulated dynamic range of their intra-axon size variability, which as a whole led to widespread thinning in lobule HV PC axons in ET.

### PC axonal pathology is linked with clinical manifestations of ET

While this study reports PC axonal pathology in the white matter (WM) compartment of the cerebellum, previous research has documented PC pathology in ET and spinocerebellar ataxias across the proximal axon segment (e.g. torpedo), soma (e.g. cell loss), and dendrites (e.g. dendritic swelling) (24, 39–41) (**Fig. 4C**). Here, we analyzed how WM PC axonal measures correlate with (1) ET severity and with (2) other pathological PC measures. Seven WM PC metrics were analyzed: HV CSA, HV CV, HV intraCV, HV intraSD, HV intraCVCV, Cr1 CSA, and Cr1 CV.

**Figure 4.**
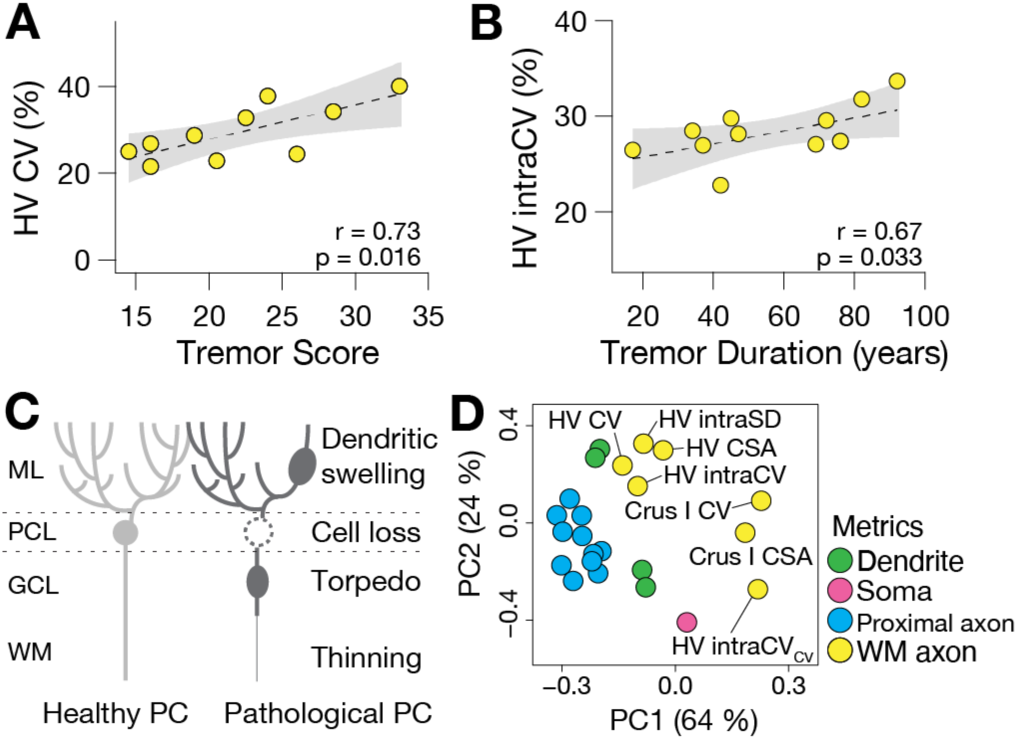
Relationship between WM PC axonal measurements and ET manifestations. **A.** Correlation between tremor score and HV CV. Each point is individual patient analyzed in this study (n = 10). Significant positive correlation is shown (Pearson’s r = 0.73, *p* = 0.016). Shaded area indicates 95% confidence interval. **B.** Correlation between tremor duration and HV intraCV. Similar format to A. Significant positive correlation is shown (Pearson’s r = 0.67, *p* = 0.033). **C.** Schematic examples of representative PC pathology in its multiple cellular components in the cerebellar cortex and white matter. Dendritic swelling, cell body loss, axonal torpedo, and axonal thinning are illustrated. **D.** PCA plot separating white matter and other PC pathological measures in 10 ET patients. Each point represents a metric (e.g., HV CV). Analysis includes dendritic (green), somatic (red), and proximal axonal (blue) metrics that have previously been established as significant contributors to disease manifestation in a study of the same ET patients (n=19 metrics) (31), as well as WM axon metrics presented in the current analyses (yellow). Complete list of PC pathological metrics and their intercorrelations are shown in Fig. S1. Abbreviations: CSA, cross sectional area; GCL, granule cell layer; ML, molecular layer; PCL, Purkinje cell layer; WM, white matter.

First, using the clinical history of the patients, we quantified correlations between tremor scores and WM PC axonal morphology. Among these, HV CV and tremor score resulted in the strongest correlation (r = 0.73, *p* = 0.016; **Fig. 4A**). HV CSA, HV intraSD, and HV intraCVCV were not significantly correlated with the tremor score (r = 0.02, 0.20, and 0.36; *p* = 0.95, 0.58, and 0.30; respectively), despite their significant changes in ET (**Figs. 3F,M,N**). The other metrics including HV intraCV, Cr1 CSA, and Cr1 CV also did not correlate with tremor score. Correlations between tremor duration and WM PC axonal morphology were similarly quantified. In this comparison, HV intraCV showed the strongest correlation (r = 0.67, *p* = 0.033; **Fig. 4B**), while the other metrics did not show significant correlations. In summary, while there were some intriguing clinical-pathological correlations that were restricted to HV, these were not linked to the measures that differentiated ET patients from controls.

The present study utilized subjects who were previously analyzed to find significant correlations between clinical manifestations and multiple PC metrics (4 dendritic, 1 somatic, and 14 proximal axonal measures (24, 29–31); **Fig. S1**). To determine whether measures of the PC WM compartment could reveal an unrecognized source of variability in ET patients, we performed principal component analysis (PCA) on combined data from current and previous studies (n = 10 ET patients). The resulting PCA plot revealed that metrics describing WM PC axon morphology occupied a space separate from those of other PC pathological metrics (**Fig. 4D**). This indicated that while all measures correlate independently with disease manifestation, they may represent qualitatively distinct sets of features within these patients. The weak correlations between these metric groups (e.g. dendrite, soma, etc.) support their qualitative differences (**Fig. S1**).

Together, these analyses highlight PC axonal changes in the cerebellar WM as potentially unique metrics of cerebellar degeneration in ET, with some intriguing clinical-pathological correlations that deserve additional attention in future studies with larger sample sizes.

## Discussion

Growing evidence points to differential disease susceptibility among known PC subtypes in animal models, but the relevance of these findings to human cerebellar pathology were unknown. Here, we analyzed PC axonal pathology in folial white matter in the post-mortem cerebellar cortex of individuals with ET, a prevalent neurological disorder. In control subjects, we first identified two distinct PC populations in motor lobule HV and cognitive lobule Crus I on the basis of different NEFH expression levels and axon caliber. In ET patients, we found PC axonal atrophy selectively in HV folial white matter, while axons in Crus I white matter maintained normal morphology. Intriguing clinical-pathological correlations were also evident, although these should be regarded as preliminary. This region- and cell type-specific pathology provides the first evidence of PC subtype vulnerability in the human cerebellum. The relevance of these findings to cerebellar pathophysiology are discussed.

### PC heterogeneity in human cerebellum

Cerebellar lobules contain specific combinations of distinct sets of PCs (3, 32). In rodents, PCs are broadly categorized into two major groups based on positive or negative Aldolase C (Aldoc) expression. These groups differ in their input-output circuits, spiking properties, modes of plasticity, and gene expression profiles (33–39). In primates, however, Aldoc expression patterns become non-binary, complicating PC subtype identification (40–42). To probe the organization of the human cerebellum, we therefore examined PCs using NEFH expression and axon caliber, markers that have been shown to distinguish Aldoc subtypes (26, 43). Specifically, we focused on lobules HV and Crus I, since they represent two extremes of the functional gradient between cerebellar motor and cognitive function, which broadly correspond with the distribution of Aldoc- and Aldoc+ PCs, respectively (10, 39, 44). Here, we show that human HV and Crus I contain NEFH-high PCs with thick axons and NEFH-low PCs with thinner axons, respectively (**Fig. 2**). Because NEFH expression is also known to correlate with ion channel composition, spiking capacity, and synaptic properties (45), our data indicate that the signals at HV and Crus I are processed by anatomically and physiologically distinct PC populations. Considering the widely conserved connectivity pattern of lobules HV and Crus I with dorsal and ventral cerebellar dentate nucleus (19, 46), respectively, and with segregated downstream circuits (7), our data further predict that the lobule-specific disruption of PCs in either lobule could result in different functional consequences in humans (8, 11, 33, 47–50).

PC subtypes have long been thought to differ in their susceptibility to disease, as evidenced by several disease models that exhibit patterned PC loss. For instance, in animal models of hypoxia, Nieman-Pick type C, spinocerebellar ataxia type 1 (SCA1), and autosomal recessive spastic ataxia of Charlevoix-Saguenay, PC loss is most pronounced in Aldoc- stripes of the cerebellar cortex (1, 51–54). In humans, patterned cerebellar atrophy has been reported in the SCAs, aging, and alcoholism (55–57), though studies to date lack the PC cell type resolution that helps to understand circuit connectivity and functional consequences. Using ET post-mortem cerebella, we demonstrate region- and cell type-specific PC vulnerability patterns in humans. PCs in lobule HV, which processes upper limb and hand movement-related information (58), were enriched in structurally atrophic axons (**Fig. 3**). These PCs are molecularly distinct from those in lobule Crus I, the primary cognitive region of the cerebellum, where axons in ET were structurally indistinguishable from control. Together, the results underscore the importance of PC subtype characterization in understanding cerebellar degeneration.

### Implications for pathophysiology of essential tremor

ET is among the most common neurodegenerative disorders globally and is characterized by an 8-12 Hz kinetic tremor of the upper limbs (20, 22, 59, 60). Classically, ET pathophysiology was hypothesized to originate in the inferior olive, whose collateralizing inputs to PCs fire at a frequency similar to that of the tremor in ET (61). These inputs were thought to synchronize PC activity and facilitate time-locked cerebellar motor outputs (25, 62, 63). In contrast, more recent post-mortem studies have identified degenerative changes centered in PCs, indicating that the cerebellar cortex also plays a major role in disease pathophysiology (25, 31). To further contextualize the role of the cerebellum in the generation of motor symptoms, we provide novel metrics of axonal pathology, at both individual and population levels. The WM PC axonal morphology metrics add to the array of other well-characterized cerebellar cortical measures of ET pathology and are also predictive of ET pathophysiology independent from these cortical measures (**Fig. 4**).

Separation of WM axonal pathology from the proximal axonal pathology is further supported by our observations that PCs with proximal axonal pathology (e.g. torpedo) often have a recurrent collateral process and lack clear axonal extension into WM (30, 31).

Axon caliber is a well-known determinant of electrical conduction velocity and fidelity (64, 65), and is tightly correlated with expression levels of neurofilament molecules (12, 13, 17, 66). In ET, we showed that HV PC axons were not only thinner but exhibited decreased intra-axon variable range and dysregulation of this range across axons (**Fig. 3K-N**).

Interestingly, this variability, but not mean size, correlated with tremor score and duration (**Fig. 4**). We conjecture that symptoms could originate from abnormal cerebellar corticonuclear signal transmission associated with a thinned and dysregulated shape of individual PC axons. Irregularity in cerebellar cortical activity, leading to irregular cerebellar nuclear responses, has been causally demonstrated in generation of tremor and dystonia; rectification of these aberrant signals has therapeutic effects (67–71). Our data thus highlights an understudied locus that contributes to the pathophysiology of movement disorders. While we do not know whether the axonal change originates in PCs themselves or the myelin sheath that supports axons, molecular processes that regulate axonal integrity will be of interest for future studies in the context of ET and movement disorders in general (18, 72–78).

### Limitations and methodological considerations

While the striped distribution of PC subsets is well-characterized (39), its detailed analysis in human cerebella remains challenging due to extensive foliation and limited post-mortem tissue availability (79). Therefore, we focused our analysis on broader PC subtype differences between lobules(32), using axonal morphology as cross-validation for PC subtype identification.

To analyze axonal morphology, we have chosen apical regions in folial white matter for consistency. While analyses of the axons even more distal to PC somata could display distinct pathology, this would be challenging because deeper white matter is mixed with PC axons from the lobular apex and base, containing both distal and proximal axons, respectively, and the white matter surrounding the cerebellar nuclei are indistinguishable with respect to their lobules of origin. Axonal caliber measures were gathered with 3D reconstruction of confocal images as opposed to electron microscopy, which is considered a standard for this analysis. Standard transmission electron microscopy of human brain autopsy material has limited application in neurobiological research due to fixation and autolysis issues.

## Conclusion

We identified that human motor lobule HV and cognitive lobule Crus I consist of discrete sets of PCs with distinct axonal caliber and NEFH expression. In the ET cerebellum, HV PCs demonstrated thinning in the WM, with a greater degree of across- and intra-axonal caliber variability that correlate with clinical metrics, whereas Crus I WM PC axons were unchanged in ET. These results demonstrate the first example of selective vulnerability of specific PC subtypes to neurodegenerative disease in the human cerebellum.

## Methods

### Ethics statements

Analyses included 18 brains in total. All study subjects signed informed consent forms approved by the respective university or institutional ethics boards.

### Post-mortem cerebellum acquisition, neuropathological and clinical evaluation

Formalin-fixed cerebellar samples from 12 ET patients and 6 controls (2:1 matching) were obtained from the New York Brain Bank at Columbia University. ET brains were acquired through the Essential Tremor Centralized Brain Repository (ETCBR), a collaboration between investigators at UT Southwestern Medical Center and Columbia University Medical Center. Case selection was guided by the availability of recently obtained cerebellar parasagittal tissue slices containing both HV and Crus I. Two ET subjects with low immunoreactivity for PCP2 were excluded from the analysis, resulting in a final sample of 10 ET patients.

ET donors were followed prospectively, and diagnoses were made by an experienced movement disorders neurologist (EDL) using three sequential methods, as previously described (80, 81). Using a standardized, videotaped neurological assessment, EDL assessed postural, kinetic, intention, and rest tremors. Action tremor was rated in each arm using a series of 12 tasks (score = 0 – 18 for each arm), yielding a total tremor score (score = 0 – 36). The presence of other movement disorders (e.g. dystonia, Parkinsonian features) were also assessed in detail. Patients with a history of traumatic brain injury, excessive alcohol consumption, and/or use of medications known to cause cerebellar damage were excluded from all analyses in this study (80). Control donors were acquired through the Alzheimer’s Disease Research Center or the Washington Heights Inwood Columbia Aging Project, Columbia University. During serial neurological examinations, these individuals were found to be clinically free of ET and other neurodegenerative disorders, including Alzheimer’s disease, Parkinson’s disease, or progressive supranuclear palsy.

All patients included in these studies agreed to and signed informed consent forms approved by the respective university ethics boards overseeing their participation. At the time of death, brains were harvested and banked at the New York Brain Bank, where they underwent a rigorous neuropathological evaluation. Eighteen standardized blocks were harvested from whole brains and 7 μm thick paraffin sections were stained with Luxol fast blue/hematoxylin and eosin (LH&E). Selected sections were stained by immunohistochemistry for alpha-synuclein, beta-amyloid, and hyperphosphorylated tau in addition to Bielschowsky silver staining, as previously detailed (82, 83). Braak and Braak Alzheimer’s disease neurofibrillary tangle staging (84) as well as Consortium to Establish a Registry for Alzheimer’s Disease (CERAD) ratings for neuritic plaques were assigned to each brain (85). Post-mortem interval (time between patient death and brain samples freezing) and total brain weight were recorded.

### Tissue processing and immunohistochemistry

The cerebellum was dissected into 1.0 – 1.5 cm thick parasagittal slices and stored in 10% formalin at room temperature. A section at or just lateral to the dentate nucleus was used in the present study, located 1.5 – 2.5 cm from the cerebellar midline. Cerebellar lobules were identified by taking photos of parasagittal slices using a stereoscope (Stemi 508, Zeiss) and referring to a cerebellar atlas (86). We dissected an approximately 1 cm x 1 cm region at the apices of HV and Crus I (**Fig. 1**). The dissected blocks were then soaked in phosphate buffered saline (PBS, pH 7.4) containing 30% sucrose overnight at 4 °C. Subsequently, the brain blocks were embedded in gelatin by placing them into a hot gelatin solution (12% gelatin and 10% sucrose in PBS) immediately followed by cooling on ice to harden the gelatin. The hardened gelatin blocks were fixed overnight in 4% PFA, 30% sucrose in PBS. Excess gelatin was trimmed to create a ∼1 mm border around the tissue.

The gelatin blocks containing the cerebellar tissue were cut into serial sections at a thickness of 80 µm with a freezing microtome (HM430, Microm [currently Epredia, Kalamazoo, MI]). In a few cases whose parasagittal blocks were especially thin and contained limited intact lobule HV and Crus I, we sectioned at 40 µm. The tissue sections were rinsed with PBS and stored in PBS containing ∼1% sodium azide at 4 °C until use. In all cases, 3D-reconstruction analyses occurred over a depth of 30 µm.

All immunostaining experiments utilized slide-mounted tissue. Sections were mounted on TruBond 380 adhesive slides (#63700-W1, Electron Microscopy Sciences, Hatfield, PA) and dried at 30 °C on a heating plate for 30 minutes. To perform heat-induced antigen retrieval, slides were immersed in a tris-based antigen unmasking solution (#H-3301, Vector Laboratories, Newark, CA) and heated for 40 – 60 minutes in a commercial vegetable steamer. After washing in PBS, a hydrophobic boundary was drawn around individual sections using an ImmEdge Pen (#H-4000, Vector Laboratories). The sections were incubated for 1 hour at room temperature in blocking buffer (10% normal donkey serum and 1% bovine serum albumin in PBS containing 0.1 % Triton-X (PBST)), then incubated in blocking buffer containing primary antibody at 4 °C overnight. The following primary antibodies were used. For axon reconstruction studies, we used a mouse anti-PCP2 monoclonal antibody at a concentration of 0.2 µg/ml (#sc-137064, Santa Cruz Biotechnology, Dallas, TX). To measure PC neurofilament expression, a mouse monoclonal antibody raised against nonphosphorylated neurofilament heavy chain was used at a concentration of 2 µg/ml (a.k.a. SMI-32; #801701, BioLegend, San Diego, CA). For PC counts, a mouse anti-Calbindin antibody at a concentration of 5-10 µg/ml (#C9848, Sigma-Aldrich, Saint Louis, MO) was used. The following day, sections were washed and then incubated in blocking buffer containing secondary antibody, donkey anti-mouse IgG antibody conjugated to Alexa Fluor 594 (∼3.75 µg/ml, made by 1:400 dilution, #715-586- 150, Jackson ImmunoResearch, West Grove, PA), in blocking buffer for 4 hours at room temperature. Washes occurred between reagents and were performed using PBS, except for the final wash, in which we used deionized water. After washing, sections were incubated in TrueBlack (#23007, Biotium, Fremont, CA) diluted 30-fold in 70% EtOH for 30 seconds on a shaker plate at low speed at room temperature to quench lipofuscin autofluorescence. Finally, slides were washed in PBS, coverslipped with Prolong Glass Antifade Mountant (#P36984, Invitrogen, Waltham, MA), and held in the dark overnight at room temperature before being stored at -20°C.

### Confocal imaging and PC axon 3D reconstruction

For PC axon reconstruction, PC axons were selectively visualized using PCP2 instead of calbindin since PCP2 is exclusively present in PC axons while calbindin is also present in olivocerebellar climbing fibers as well and could lead to a misrepresentation of PC axon caliber (87, 88). Immunostained PC axonal tracts from all samples were imaged using a LSM880 or LSM980 confocal microscope (Zeiss, Oberkochen, Germany).

To make comparison of PC axons across cases, images were taken at a consistent location: an area slightly basal to the zone where white matter from two apical folia converges (**Fig. 1**). Within this location, some variability was inevitable due to individual differences in cerebellar morphology, plane of sectioning, and slight differences in the distance of the starting parasagittal block from the cerebellar midline; however, effort was made to maintain as much consistency as possible.

Z-stacks were acquired in Zen (Zeiss) using a 63X/NA1.4 or 40X/NA1.4 oil objective. From a preliminary analysis of 40X epifluorescence images in a control sample, the thinnest group of PC axons in the cerebellar regions of interest had an average 2D diameter of 0.8 µm.

Thus, by Nyquist theorem, z-stacks were captured using a step size of 0.4 µm, covering a total depth of 30 ± 5µm, to accurately represent the entirety of the axon in all dimensions. Confocal Z-stacks were imported into Fiji, where brightness and contrast were adjusted and outliers were removed (radius = 2.0 pixels, threshold = 50). The boundary of the axon’s fluorescent signal remained the same during this process. These image sequences were saved as TIFF files and imported into Imaris 10.1.0 (Oxford Instruments, Abingdon, United Kingdom), where we utilized the “filament tracer” module to 3D-reconstruct PC axonal segments. Using the “autodepth tracing” method with automatic centering and automatic diameter options enabled, trained laboratory personnel manually traced all axon segments, after which Imaris created 3D renderings with its intensity thresholding algorithm. Following 3D reconstruction, each segment was manually examined for accuracy of reconstruction. Any axons that were poorly reconstructed (i.e., the 3D render did not accurately represent the intensity profile of the axon), were excluded from the dataset. At times, one portion of an axon was accurately reconstructed, while another portion of that same axon was not. In those cases, the inaccurate portion was removed from the axon such that only the accurate portion would remain to represent that axon’s metrics. The volume for each individual axon was divided by its length to obtain an average CSA for that axon. The CSAs of all axons from each control or ET sample were averaged to obtain a single data point representing all individual axons. Sample means were then averaged to create control or ET means. To represent the distribution of all analyzed axons, CSA values of individual axons were also converted to diameters assuming their circularity.

### Measuring intra-axon variability of 3D PC axon profiles

To represent the variability of individual axons along their length, we analyzed the profiles of our 3D-reconstructed axons in Rhinoceros 7 (Robert McNeel & Associates, Seattle, WA), a 3D modeling software. 3D reconstructions from Imaris were exported as WRL files for import into Rhinoceros. We wrote custom code using Rhinoceros’ native Python library that extracted cross-sectional areas at 0.5 µm intervals along the profiles of individual axons. From these values, a standard deviation and coefficient of variation (%) were calculated for each axon to represent its intra-axonal variability. Measures of Intra-axonal variability from all axons analyzed for a single ET case or control were averaged into a single data point to represent that subject.

### Quantification of neurofilament expression in PC

Expression levels of NEFH in the PCs were quantified from fluorescently immunostained sections and were compared between lobules HV and Crus I. The sections adjacent to those used for axon reconstructions were used for this analysis. To minimize the variability of staining between the compared regions, pairs of HV and Crus I sections from the same individuals were simultaneously stained on the same slides. All images were captured using identical exposure time and intensity. Images for analysis were captured using an epi-fluorescence microscope, Zeiss Axio Imager.M2 with ORCA-Flash4.0LT Plus CMOS camera (Hamamatsu Photonics, Hamamatsu, Japan) and 10X/NA0.3 lens. In reliably stained regions from each section, PCs whose somata did not overlap with neighboring PCs and whose nuclei were easily identified were quantified. In ImageJ, pixel intensity (0 – 255 a.u.) was obtained within manually circumscribed contours of individual PC somata. For group comparison, mean intensities for each PC were averaged into sample means and compared across lobules.

### Quantification of the fraction of PCP2-expressing PCs

The fraction of PCP2-expressing PCs was quantified in the cerebellar sections fluorescently immunostained for PCP2 with Nissl counterstaining. During immunostaining for PCP2, a 1:200 dilution of NeuroTrace 435/455 (N21749, ThermoFisher, Waltham, MA) was added to the blocking buffer containing secondary antibody and incubated for 4 hours at room temperature. Subsequently, two-color images were taken from the apical-most folia of the lobules HV and Crus I using an epi-fluorescent microscope with 10X/NA0.3 lens. The PCs were manually identified by their location in the PC layer, large somata, and by the presence of a soma and nucleus in the channel for Nissl counterstaining. These PCs were then cross-referenced to the PCP2 channel to determine co-labeling. Counts of PCP2 only, Nissl only, or co-expression were manually obtained.

### Statistics and data analysis

All analyses were conducted by lab personnel who were blinded to all clinical information and diagnosis. Statistical analyses were performed and visualized in Prism 10 (GraphPad Software, Boston, MA) Igor Pro 9 (WaveMetrics, Portland, OR), or R (v4.0.5; https://www.r-project.org/). All values were reported as mean ± standard deviation, unless stated otherwise. Raw data were subjected to Kolmogorov-Smirnov normality testing, after which appropriate statistical tests were determined. Clinical findings were compared using an unpaired t-test, Fisher’s exact test, or a chi-square test, as applicable. Within-subject comparisons of post-mortem metrics (e.g. axon caliber, NEFH expression, etc.) in control samples utilized paired t-tests since data were all normally distributed. Case-control comparisons of post-mortem metrics utilized unpaired t-tests with Welch’s correction since sample sizes differed. Linear regression analyses were used to compare relationships between normally distributed variables and *p* values were calculated to represent whether the slope was significantly non-zero. Pearson’s correlation coefficients are reported as ± r. For data analysis in Figure 4, we compared previously published cerebellar pathology data (24, 29–31) with our new white matter Purkinje cell (PC) axonal measurements within the same 10 ET patients. Using principal component analysis in R, we examined 7 WM axonal, 4 dendritic, 1 somatic, and 14 proximal axonal PC pathology metrics (see **Fig. S1** for specific metrics). We analyzed their associations using Z-scores. Missing data points were excluded when calculating the covariance matrix.

## Acknowledgements

This work was supported by National Ataxia Foundation (HF), SWMF/Kriscunus Tremor Funding (EDL), NIH R01 NS086736 (PLF and EDL), and NIH R01 NS117745 (EDL). The authors would like to acknowledge the Quantitative Light Microscopy Core, a Shared Resource of the Harold C. Simmons Cancer Center, supported in part by an NCI Cancer Center Support Grant, 1P30 CA142543-01. Use of LSM880/980 microscopes was supported by NIH 1S10 OD021684-01 to Kate Luby-Phelps (UT Southwestern). We thank all the patients and families that contributed to brain donation. We thank Juliana Neniel for assistance in editing the final version of the paper.

## Author contributions

Conceptualization, JW, EDL, HF; Methodology, JW, HF; Software, JW, HF; Validation, JW, PLF, EDL, HF; Formal analysis, JW, HF; Investigation, JW, HF; Resources, PLF, EDL, HF; Data Curation, JW, PLF, EDL, HF; Writing -Original Draft, JW, HF; Writing - Review & Editing, JW, PLF, EDL, HF; Visualization, JW, HF; Supervision, EDL, HF; Funding acquisition, PLF, EDL, HF.

## Competing interest statement

We do not have any competing interests.

## Notes

### Competing Interest Statement

The authors have declared no competing interest.

